# SHRIMP: Genetically Encoded mScarlet-derived Red Fluorescent Hydrogen Peroxide Sensor with High Brightness and Minimal Photoactivation

**DOI:** 10.1101/2023.08.09.552302

**Authors:** Yu Pang, Yiyu Zhang, Jing Zhang, Zefan Li, Yi He, Yong Wang, Jose Oberholzer, Hui-wang Ai

**Author notes:** Corresponding author: Hui-wang Ai., **Email:**. These two authors contributed to this work equally (Yu Pang, Yiyu Zhang).

## Abstract

Red fluorescent protein (RFP) derived indicators are popular due to advantages such as increased imaging depth and reduced autofluorescence and cytotoxicity. However, most RFP-based indicators have low brightness and are susceptible to blue-light-induced photoactivation. In this study, we aimed to overcome the limitations of existing red fluorescent indicators. We utilized mScarlet-I, a highly bright and robust monomeric RFP, to develop a circularly permuted variant called cpmScarlet. We further engineered cpmScarlet into a novel red fluorescent indicator specifically for hydrogen peroxide (H_2_O_2_), a crucial reactive oxygen species (ROS) involved in redox signaling and oxidative stress. The resultant indicator, SHRIMP (mScarlet-derived H_2_O_2_ Redox Indicator with Minimal Photoactivation), exhibited excitation and emission peaks at ∼570 and 595 nm, respectively, and demonstrated a maximum five-fold fluorescence turn-off response to H_2_O_2_. Importantly, SHRIMP was not susceptible to blue-light-induced photoactivation and showed high brightness both in its purified protein form and when expressed in mammalian cells. We successfully employed SHRIMP to visualize H_2_O_2_ dynamics in mammalian cells with exogenously added H_2_O_2_ and in activated macrophages. Additionally, we demonstrated its utility for multiparameter imaging by co-expressing SHRIMP with GCaMP6m, a green fluorescent calcium indicator, enabling simultaneous monitoring of H_2_O_2_ and calcium dynamics in mammalian cells in response to thapsigargin (TG) and epidermal growth factor (EGF) stimulation. Furthermore, we expressed SHRIMP in isolated primary mouse islet tissue, and SHRIMP exhibited excellent brightness and capability for effective detection of H_2_O_2_ production during streptozotocin (STZ)-induced β-cell damage. This study successfully transformed mScarlet-I, a bright and robust monomeric RFP, into a circularly permuted variant (cpmScarlet) and developed the first cpmScarlet-based genetically encoded fluorescent indicator called SHRIMP. SHRIMP exhibits high brightness and insensitivity to photoactivation and is a valuable tool for real-time monitoring of H_2_O_2_ dynamics in various biological systems. Further research may yield an expanded family of cpmScarlet-based red fluorescent indicators with enhanced photophysical properties.

## Introduction

Genetically encoded fluorescent indicators (GEFIs) are indispensable for studying real-time dynamics and activities in biological systems (1,2). While green fluorescent proteins (GFPs) and yellow fluorescent proteins (YFPs) have been widely used for developing successful single fluorescent protein (FP)-based GEFIs, there is growing interest in red fluorescent GEFIs due to their potential for deeper tissue penetration, reduced cytotoxicity, and lower background fluorescence (3,4). Red GEFIs can also be combined with green GEFIs for multi-color imaging and simultaneous monitoring of multiple biological processes.

Circular permutation of FPs is a common strategy for developing GEFIs by fusing sensory domains to the FP (5,6). The circular permutation process creates new N- and C-termini near the FP chromophore, allowing the fusion of sensory domains to modulate FP fluorescence. In previous studies, circularly permuted variants of RFPs such as mApple, mRuby, mCherry, and FusionRed have been engineered into GEFIs that can sense calcium ion (Ca^2+^) or other biological parameters (7-15). However, red GEFIs, especially when expressed in cells, tend to be less bright than their green counterparts, and engineering responsive red GEFIs is generally more challenging (3,16). Furthermore, blue-light-induced photoactivation has been observed for mApple and cpmApple-derived GEFIs (8,10,17), potentially leading to artifacts and data interpretation issues. The problems of existing red GEFIs are reasoned to be inherited from their parental RFPs, which are less bright and less robust than GFPs and YFPs (17).

Redox signaling plays a crucial role in regulating various cellular processes, including cell growth, differentiation, and apoptosis (14,18). When the balance between reactive oxygen species (ROS) production and antioxidant defense mechanisms is disrupted, it can lead to oxidative stress, which is characterized by an excess of ROS that can cause damage to cellular macromolecules such as proteins, lipids, and DNA. Oxidative stress has been implicated in the pathogenesis of many diseases, including cancer, cardiovascular disease, and neurodegenerative disorders (19-22). In this context, GEFIs that can sense ROS and redox dynamics are useful tools not only for mechanistic studies but also for developing effective therapies for diseases (14,23,24). Several red fluorescent redox indicators have been developed in previous studies, including some by our team (12,14,25-29).

Hydrogen peroxide (H_2_O_2_) is a key reactive oxygen species (ROS) generated from the incomplete reduction of oxygen (30,31). It is a relatively stable ROS that can diffuse within or between cells. H_2_O_2_ mediates redox signaling by oxidizing redox-sensitive cysteine residues in proteins, leading to subsequent conformational and activity changes (32,33). It has been identified as a second messenger involved in insulin and growth factor-induced signaling and protein kinase regulation (32,34-36). However, high levels of H_2_O_2_ can have irreversible and often harmful effects, making it a toxic metabolite (37,38). Previous work has generated a family of GEFIs for H_2_O_2_, and the most notable examples are HyPer variants based on a circularly permuted YFP (39-41). The only available red GEFI for H_2_O_2_, HyPerRed (25), was generated by inserting cpmApple into the regulatory domain of a bacterial H_2_O_2_-sensing protein OxyR.

In our attempts to use cpmApple-derived redox indicators for tissue imaging, we faced challenges in obtaining consistent results. To address this, we turned to mScarlet (42), a recently reported monomeric RFP with remarkable brightness, folding robustness, fast chromophore maturation, and resistance to blue-light-induced photoactivation. In this study, we present our progress in engineering a circularly permuted variant of mScarlet (cpmScarlet) and the further development and validation of SHRIMP, a novel mScarlet-based H_2_O_2_ redox indicator with high brightness and minimal photoactivation. We successfully used SHRIMP to image real-time H_2_O_2_ dynamics in mammalian cell lines and pancreatic islet tissue. Our findings highlight the potential of cpmScarlet as a starting point for engineering improved red GEFIs and demonstrate the utility of SHRIMP for investigating H_2_O_2_ dynamics in living systems.

## Results

### Development of a Circularly Permuted mScarlet Variant

When we started this project, mScarlet was reported as a new RFP exhibiting exceptional brightness and quantum yield (42). Among its variants, mScarlet-I stood out with its favorable combination of a reasonable quantum yield (0.54), rapid maturation, and high brightness in live cells. Notably, compared to other tested RFPs, mScarlet-I displayed the highest sensitized emission when used as a Förster resonance energy transfer (FRET) acceptor for YFP (42). Considering these characteristics, we selected mScarlet-I as the foundation for our study.

Our goal was to circularly permute mScarlet-I at β-strand 7 of the FP fold, a common site for circular permutation due to its proximity to the chromophore (6). We genetically amplified two fragments of mScarlet-I, residues 149-232 and residues 2-146, and connected them using a Gly-Ser-rich linker. Additional randomized residues were introduced at the new N- and C-termini, with guidance from our recently reported enhanced circular permuted mApple (ecpApple) variant (29,43). The resulting gene library was introduced into *E. coli*, and colonies were screened for fast fluorescence onset after overnight incubation at 37 °C. Promising colonies were selected for liquid culture inoculation. After inducing protein expression and preparing cell lysates, brightness screening using a microplate reader identified the brightest clone, which we named cpmScarlet. We compared cpmScarlet with the ecpApple variant and found that cpmScarlet exhibited approximately 7.7-fold higher fluorescence in *E. coli* lysates prepared under identical conditions. Furthermore, the purified cpmScarlet protein showed approximately 6-fold higher fluorescence compared to ecpApple (**Table 1**). Its excitation peak was observed at 570 nm, and the emission peak was at 595 nm, closely resembling those of mScarlet-I (42). DNA sequencing confirmed that cpmScarlet acquired two mutations (13T and 14V) at the N-terminus and one mutation (251L) at the C-terminus.

**Table 1.** Photophysical properties of the indicated cpRFPs and red fluorescent H_2_O_2_ sensors.

| | $\lambda_{\text{ex}}$ (nm) | $\lambda_{\text{em}}$ (nm) | $\epsilon$ (M<sup>-1</sup> cm<sup>-1</sup>) | $\Phi$ | molecular brightness<sup>a</sup> |
| --- | --- | --- | --- | --- | --- |
| cpmScarlet | 570 | 595 | 31,546 | 0.31 | 9.79 |
| ecpApple | 570 | 600 | 46,924 | 0.03 | 1.60 |
| SHRIMP | reduced |  |  |  |  |
|  | 570 | 600 | 127,784 | 0.32 | 40.89 |
|  | oxidized |  |  |  |  |
|  | 570 | 600 | 25,557 | 0.32 | 8.18 |
| HyPerRed | reduced |  |  |  |  |
|  | 575 | 605 | 19,074 | 0.27 | 5.33 |
|  | oxidized |  |  |  |  |
|  | 575 | 605 | 38,148 | 0.27 | 10.30 |

<sup>a</sup> The intrinsic molecular brightness is defined as the product of $\epsilon$ and $\Phi$ .

### Engineering of the cpmScarlet-based Red Fluorescent H_2_O_2_ Indicator SHRIMP

Subsequently, we fused the fragments of the H_2_O_2_-sensing OxyR domain in HyPerRed (25) to cpmScarlet (**Fig. 1A**), ensuring that the linker length of three amino acids on each side between the OxyR fragments and cpmScarlet was preserved. To create the initial library for screening, we fully randomized the two amino acids nearest to the N- and C-termini of cpmScarlet using degenerate codons. From this library, we selected an initial mutant (named SHRIMP0.1) that exhibited approximately 2.9-fold decreased fluorescence in response to the addition of 500 μM H_2_O_2_. In addition, we identified several clones that exhibited slight fluorescence increases in response to H_2_O_2_. However, despite our further protein engineering effort, we were unable to enhance the responsiveness of these mutants.

**Figure 1.**
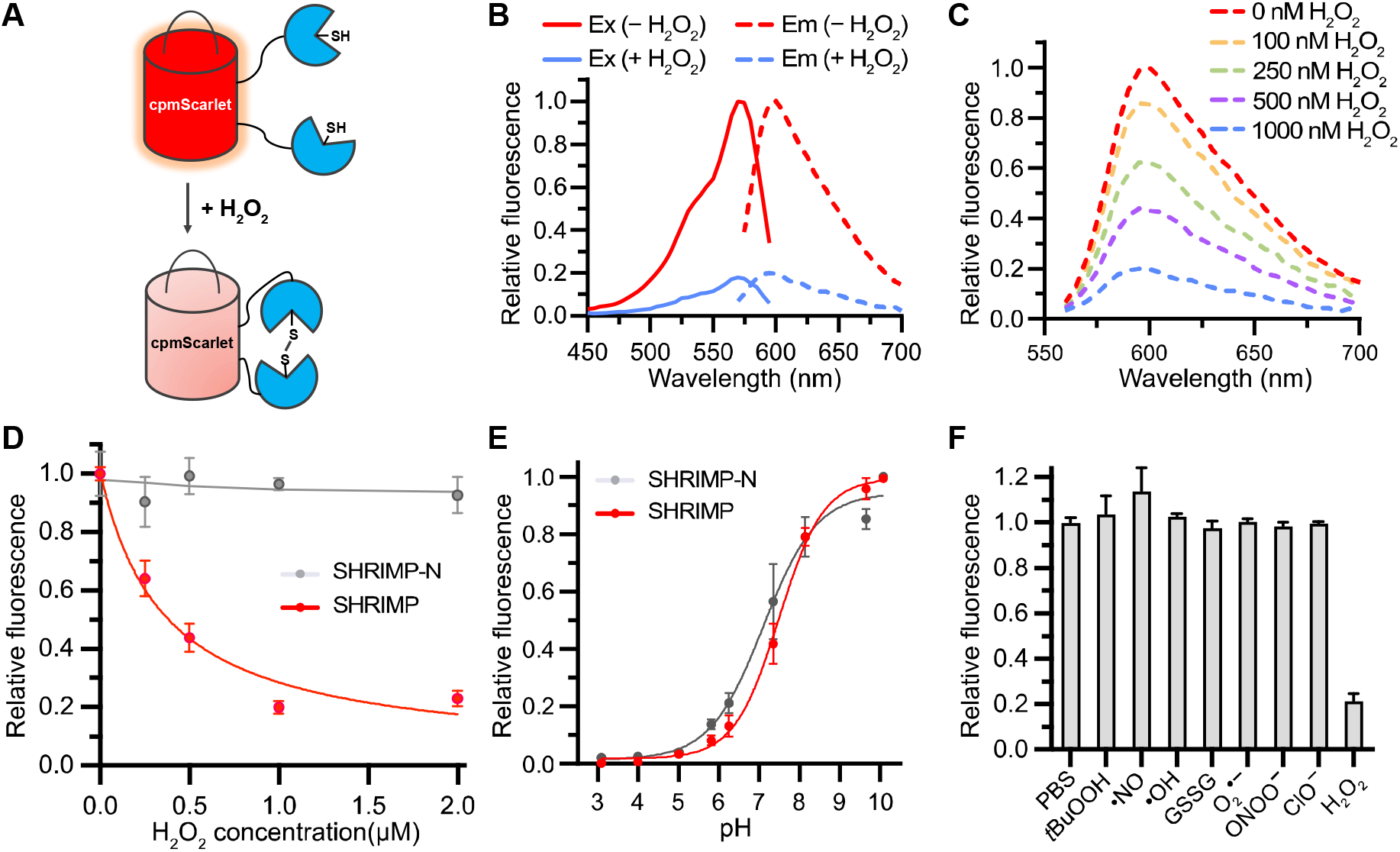
Design and *in vitro* characterization of SHRIMP. (A) Schematic illustration of the sensing mechanism of SHRIMP. (B) Excitation (solid line) and emission (dotted line) spectra of the purified SHRIMP proteins in the fully oxidized (blue) and reduced (red) states. (C) Emission spectra of SHRIMP in response to different concentrations of H_2_O_2_. (D) Fluorescence intensities of the purified SHRIMP and SHRIMP-N proteins plotted against different concentrations of H_2_O_2_. (E) pH sensitivity of SHRIMP (apparent *p*K_a_ = 7.5) and SHRIMP-N (apparent *p*K_a_ = 7.1). (F) Selectivity assay of SHRIMP among various physiologically relevant oxidants. Experiments were repeated three times and representative results are shown in panels B and C. Results in panels D-E are presented as mean and s.d. of three technical repeats.

SHRIMP 0.1 displayed relatively dim red fluorescence and required expression at 16°C. To enhance the maturation and brightness of SHRIMP 0.1, we conducted two rounds of error-prone polymerase chain reaction (PCR)-based random mutagenesis. Fluorescent colonies observed on agar plates were carefully chosen and utilized to inoculate liquid cultures in 96-well deep-well plates. For the clones derived from the first error-prone library, protein expression was induced at room temperature. Subsequently, for the clones obtained from the second-round library, we increased the expression temperature to 30 °C. In both rounds, cell lysates were prepared for each chosen clone, and their response to H_2_O_2_ was assessed using a fluorescent plate reader. The clones exhibiting the highest brightness and responsiveness to H_2_O_2_ were identified based on predefined criteria and considered as potential candidates. Among the variants obtained, one particular clone with linkers Pro-Gly-Thr and Gly-Gly-Val showed the highest fluorescence change of approximately 5-fold upon the addition of H_2_O_2_. This evolved variant gained three mutations (A363T, A404T, K495M) during the mutagenesis process. We named this clone SHRIMP (mScarlet-derived H_2_O_2_ Redox Indicator with Minimal Photoactivation) and selected it for further characterization. In addition, we created a mutant of SHRIMP that is unresponsive to H_2_O_2_ by substituting the critical cysteine residue at position 159 with serine. This mutant was named SHRIMP-N and was designed to serve as a negative control in our experiments.

### *In vitro* Characterization of SHRIMP

We expressed His_6_-tagged SHRIMP in *E. coli* and subsequently purified the protein using a combination of nickel nitrilotriacetic acid (Ni-NTA) based affinity purification and size-exclusion chromatography. We next proceeded with *in vitro* characterization of the SHRIMP protein. SHRIMP exhibited an excitation peak at 570 nm and an emission peak at 595 nm (**Fig. 1B**). Upon the addition of 1 μM H_2_O_2_, the fluorescence intensity of SHRIMP showed a maximum 5-fold decrease. The fluorescence response of SHRIMP to H_2_O_2_ followed a dose-dependent pattern, with a half-maximal response concentration (EC_50_) of approximately 294 nM (**Fig. 1CD**). As expected, SHRIMP-N showed little fluorescence response when H_2_O_2_ was added (**Fig. 1D**). To evaluate the pH sensitivity of SHRIMP and SHRIMP-N, we determined their apparent *p*K_a_ values to be 7.5 and 7.1, respectively (**Fig. 1E**). Since both variants are sensitive to pH changes around the physiological pH range, it is crucial to conduct parallel experiments using SHRIMP and SHRIMP-N to accurately distinguish fluorescence changes caused by redox alterations from those caused by pH changes. To further assess the specificity of SHRIMP, we examined its fluorescence in the presence of various oxidants, including nitric oxide (NO•), peroxynitrite (ONOO^−^), oxidized glutathione (GSSG), and other biologically relevant ROS. Notably, a drastic decrease in fluorescence intensity was observed only in the presence of H_2_O_2_ (**Fig. 1F**), confirming that SHRIMP is a specific sensor for H_2_O_2_.

### Comparison of SHRIMP with HyPerRed

We performed a comparative analysis of the photophysical properties of SHRIMP and HyPerRed, with a specific focus on brightness, photostability, and sensitivity to photoactivation. Firstly, we determined the quantum yields and extinction coefficients of purified SHRIMP and HyPerRed proteins. The quantum yield of SHRIMP remained at 0.32 in both the oxidized and reduced states, slightly higher than that of HyPerRed (0.27) in both states. The extinction coefficient of SHRIMP changed from 127,784 to 25,557 M^-1^ cm^-1^ when the fully reduced protein became fully oxidized, resulting in a five-fold fluorescence turn-off response to H_2_O_2_ (**Table 1**). In contrast, the extinction coefficient of HyPerRed changed from 19,074 to 38,148 M^-1^ cm^-1^ from the reduced to the oxidized state, resulting in a two-fold fluorescence turn-on response (**Table 1**). Consequently, the molecular brightness of SHRIMP in the brighter (reduced) state was approximately 4.0-fold higher than that of HyPerRed in the brighter (oxidized) state. Similarly, the molecular brightness of SHRIMP in the dimmer (oxidized) state was approximately 1.5-fold higher than that of HyPerRed in the dimmer (reduced) state.

To evaluate their brightness further in mammalian cells, we cloned SHRIMP and HyPerRed into a pcDNA3 mammalian expression plasmid and compared their fluorescence levels in human embryonic kidney (HEK) 293T cells 24 hours after transfection. Notably, SHRIMP-expressing cells exhibited much higher fluorescence compared to HyPerRed-expressing cells when observed under fluorescence microscopy (**Fig. 2A**). Flow cytometry analysis further confirmed a much larger population of brighter cells in the SHRIMP group (**Fig. 2A**). The average intensity of individual SHRIMP-expressing cells was more than 11-fold higher than that of HyPerRed-expressing cells. Considering that we expect SHRIMP and HyPerRed to be mostly in the reduced state in untreated HEK 293T cells, the observed brightness difference aligns with the ∼7.7-fold molecular brightness difference determined using the purified proteins (**Table 1**). The additionally higher brightness of SHRIMP in live cells suggests that it may possess superior folding and chromophore maturation.

**Figure 2.**
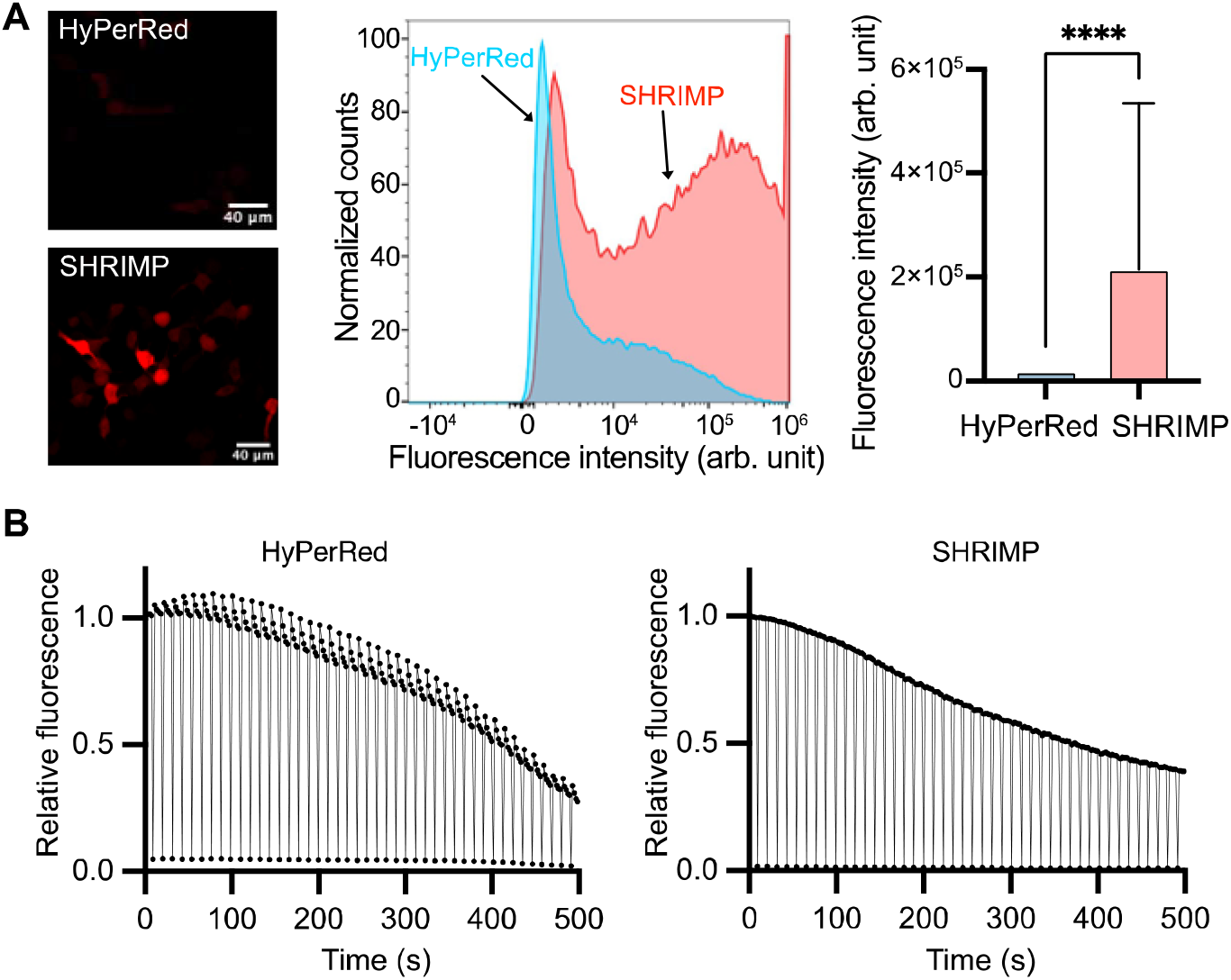
Comparison of SHRIMP and HyPerRed for mammalian cell brightness and photoactivation. (A) Comparison of the fluorescence brightness of HEK 293T cells at 24 h post transient transfection. Left: Representative fluorescence images of the cells expressing SHRIMP or HyPerRed. The images were acquired under similar conditions and presented in the same intensity scale. Middle: Flow cytometry analysis of HEK 293T cells expressing SHRIMP or HyPerRed. Right: Bar graph showing the average of individual cell fluorescence brightness from the flow cytometry experiment. The error bar represents s.d. and the statistical comparison was made with an unpaired, two-tailed t-test (\*\*\*\**P*<0.0001). (B) Fluorescence intensity traces of HyPerRed (left) and SHRIMP (right) proteins encapsulated in mineral oil droplets in response to multiple cycles of five consecutive 532 nm laser scans followed by one 405 nm laser scan on a confocal fluorescence microscope. Representative data from three independent technical repeats are presented.

To examine photostability, we subjected SHRIMP and HyPerRed proteins encapsulated in mineral oil droplets to wide-field illumination. Although HyPerRed demonstrated better photostability compared to SHRIMP under continuous high-intensity illumination (∼1.0 W/cm^2^), no significant difference in photostability between the two indicators was observed under the light intensity typically employed for time-lapse imaging experiments (∼0.091 W/cm^2^).

Violet or blue light is known to induce the photoactivation of mApple and mApple-derived GEFIs (8,10,17). We next compared the sensitivity of SHRIMP and HyPerRed to photoactivation. SHRIMP and HyPerRed proteins encapsulated in mineral oil droplets were first scanned with a 532 nm laser five times followed by a 405 nm laser scan on a confocal microscope. Multiple cycles of these scans were performed over a period of 500 s. As expected, the 532 nm laser scans led to a decrease in fluorescence for both SHRIMP and HyPerRed. The 405 nm laser scans did not photoactivate SHRIMP, but resulted in a fluorescence activation of HyPerRed ranging from 4% to 15% (**Fig. 2B**).

Taken together, our initial characterization suggests that SHRIMP has high brightness, a wide dynamic range, excellent specificity towards H_2_O_2_, reasonable photostability, and insensitivity to photoactivation.

### Imaging H_2_O_2_ Dynamic in Cultured Mammalian Cells

To further evaluate the functionality of SHRIMP in mammalian cells, we treated SHRIMP-expressing HEK 293T cells with 150 μM exogenous H_2_O_2_. SHRIMP exhibited a significant ∼52% decrease in fluorescence intensity in response to the treatment (**Fig. 3A**). In contrast, when we expressed SHRIMP-N in HEK 293T cells and subjected them to the same H_2_O_2_ treatment, no notable fluorescence change was observed (**Fig. 3A**). Considering the expected H_2_O_2_ concentration gradient of about 200-650-fold across the cell membrane (40,44), this result aligns with our *in vitro* protein assay, demonstrating that SHRIMP is responsive to nanomolar levels of H_2_O_2_.

**Figure 3.**
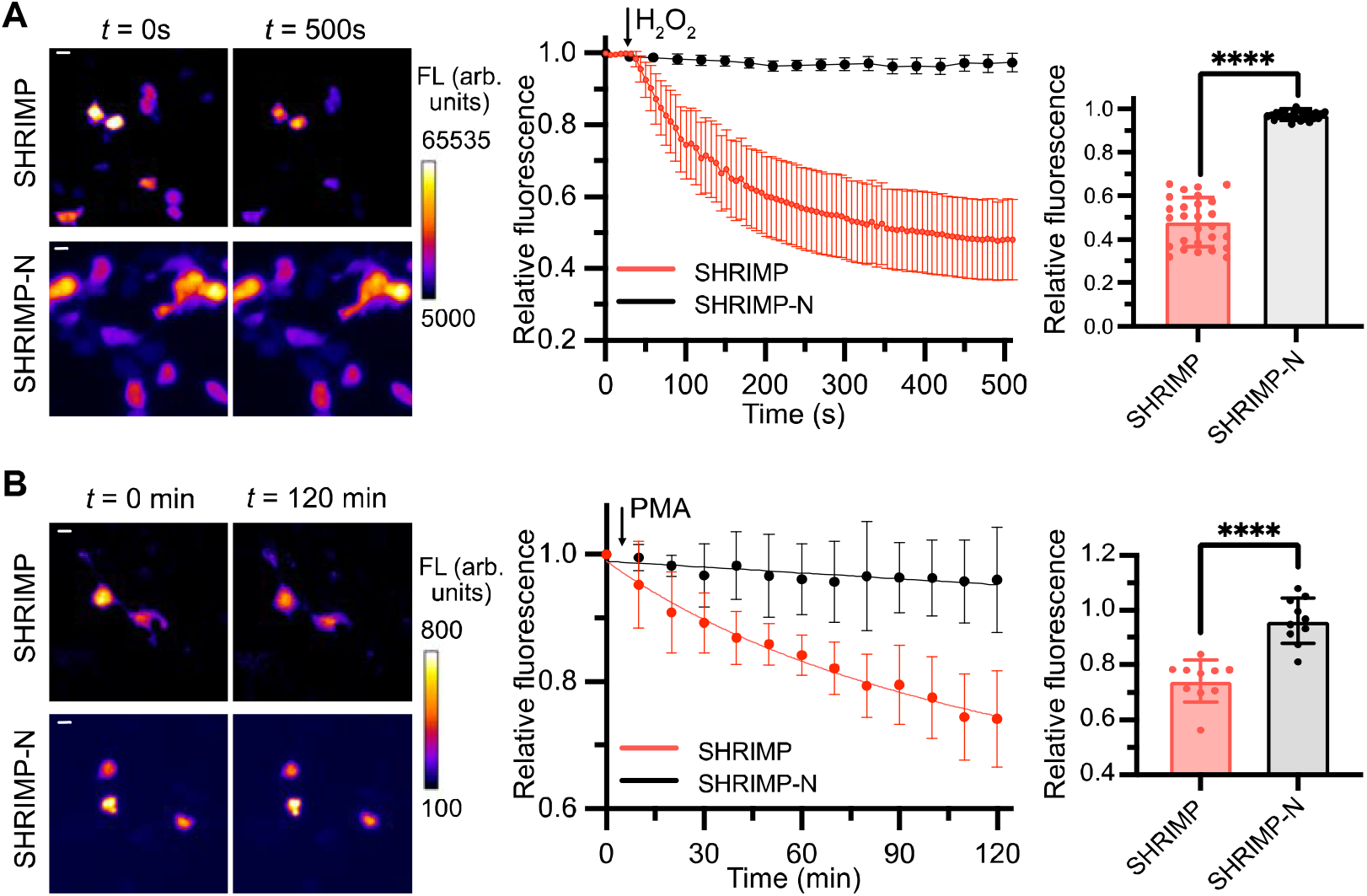
Using SHRIMP to image H_2_O_2_ in mammalian cells. (A) Fluorescence intensity changes of HEK 293T cells expressing SHRIMP or SHRIMP-N in response to extracellular H_2_O_2_ (150 μM) addition. Left: Representative pseudocolored fluorescence images. Scale bar: 10 μm. Middle: Quantitation of timelapse responses. Arrow indicates the time point for H_2_O_2_ addition. Right: Comparison of intensity changes at *t* = 500 s (****P < 0.0001, unpaired two-tailed *t* test). Data represent the mean and s.d. of 27 cells from three technical repeats in each group. (B) Fluorescence intensity changes of RAW 264.7 macrophage cells expressing SHRIMP or SHRIMP-N in response to 2 μM phorbol-12-myristate 13-acetate (PMA) activation. Left: Representative pseudocolored fluorescence images. Scale bar: 10 μm. Middle: Quantitation of timelapse responses. Arrow indicates the time point for PMA addition. Right: Comparison of intensity changes at *t* = 120 min (\*\*\*\**P*<0.0001, unpaired two-tailed t-test). Data represent the mean and s.d. of 10 cells from three technical repeats in each group. FL, fluorescence. Arb. units, arbitrary units.

To assess the applicability of SHRIMP in a physiological context, we expressed SHRIMP in mouse RAW 264.7 macrophages. It is well-established that, during phagocytosis or when activated by other stimuli, macrophages can generate superoxide and H_2_O_2_ through nicotinamide adenine dinucleotide phosphate (NADPH) oxidase and the mitochondrial respiratory chain (45). In our experiment, we transfected RAW 264.7 macrophages with SHRIMP or SHRIMP-N plasmid. After 24 hours of indicator expression, we subsequently stimulated the cells with phorbol-12-myristate 13-acetate (PMA), a protein kinase C agonist known to activate NADPH oxidase and induce H_2_O_2_ production (46,47). The RAW 264.7 cells expressing SHRIMP exhibited a notable ∼30% fluorescence decrease upon PMA stimulation (**Fig. 3B**). In parallel, we expressed SHRIMP-N in RAW 264.7 cells and subjected them to the same stimulation, there was negligible fluorescence change (**Fig. 3B**). These findings indicate that SHRIMP can monitor both exogenous H_2_O_2_ and physiological H_2_O_2_ generation in different mammalian cell systems.

### Dual-color Imaging of Ca^2+^ Signaling and H_2_O_2_ Dynamics in Live Cells

Both Ca^2+^ and H_2_O_2_ are key cellular signaling molecules, and there is substantial evidence supporting the existence of crosstalk between their signaling pathways (48-51). Consequently, there is significant interest in concurrently monitoring the dynamics of Ca^2+^ and H_2_O_2_ in live cells. To assess the compatibility of SHRIMP with green fluorescent Ca^2+^ indicators for dual-parameter imaging, we co-expressed SHRIMP in the mitochondria and GCaMP6m in the cytosol of HEK 293T cells. To confirm the proper localization of these indicators, we performed confocal imaging, which validated their respective positions within the cells (**Fig. 4A**). Subsequently, we conducted wide-field time-lapse imaging experiments and treated the cells with thapsigargin (TG), an inhibitor of sarco/endoplasmic reticulum Ca^2+^-ATPase (SERCA). TG hampers the transport of Ca^2+^ from the cytoplasm into the sarcoplasmic reticulum (SR) and endoplasmic reticulum (ER) (52,53). As anticipated, upon TG treatment, we observed a notable increase in green fluorescence intensity from GCaMP6m in the cytosol, confirming the elevation of intracellular Ca^2+^ levels (**Fig. 4B**). The gradual rise in the Ca^2+^ signal occurred through a period of ∼ 5 min and took an additional 7 min to subside Concurrently, we observed a decrease in the red fluorescence of mitochondrial SHRIMP, starting approximately 2 min after the onset of the cytosolic Ca^2+^ concentration rise. This decrease persisted throughout the monitoring period, even after the extinction of the Ca^2+^ signal (**Fig. 4C**). These findings indicate that an increase in cytosolic Ca^2+^ can induce the generation of mitochondrial H_2_O_2_, which persists for a longer duration than the Ca^2+^ signals. Furthermore, we applied H_2_O_2_ to the cells near the end of the imaging session, which caused a further decrease in red fluorescence, confirming the responsiveness of the SHRIMP sensor during our experiment (**Fig. 4C**). Interestingly, we also observed a small increase in green fluorescence upon H_2_O_2_ application, corroborating previous findings that H_2_O_2_ can reversely trigger intracellular Ca^2+^ signals (49,54,55).

**Figure 4.**
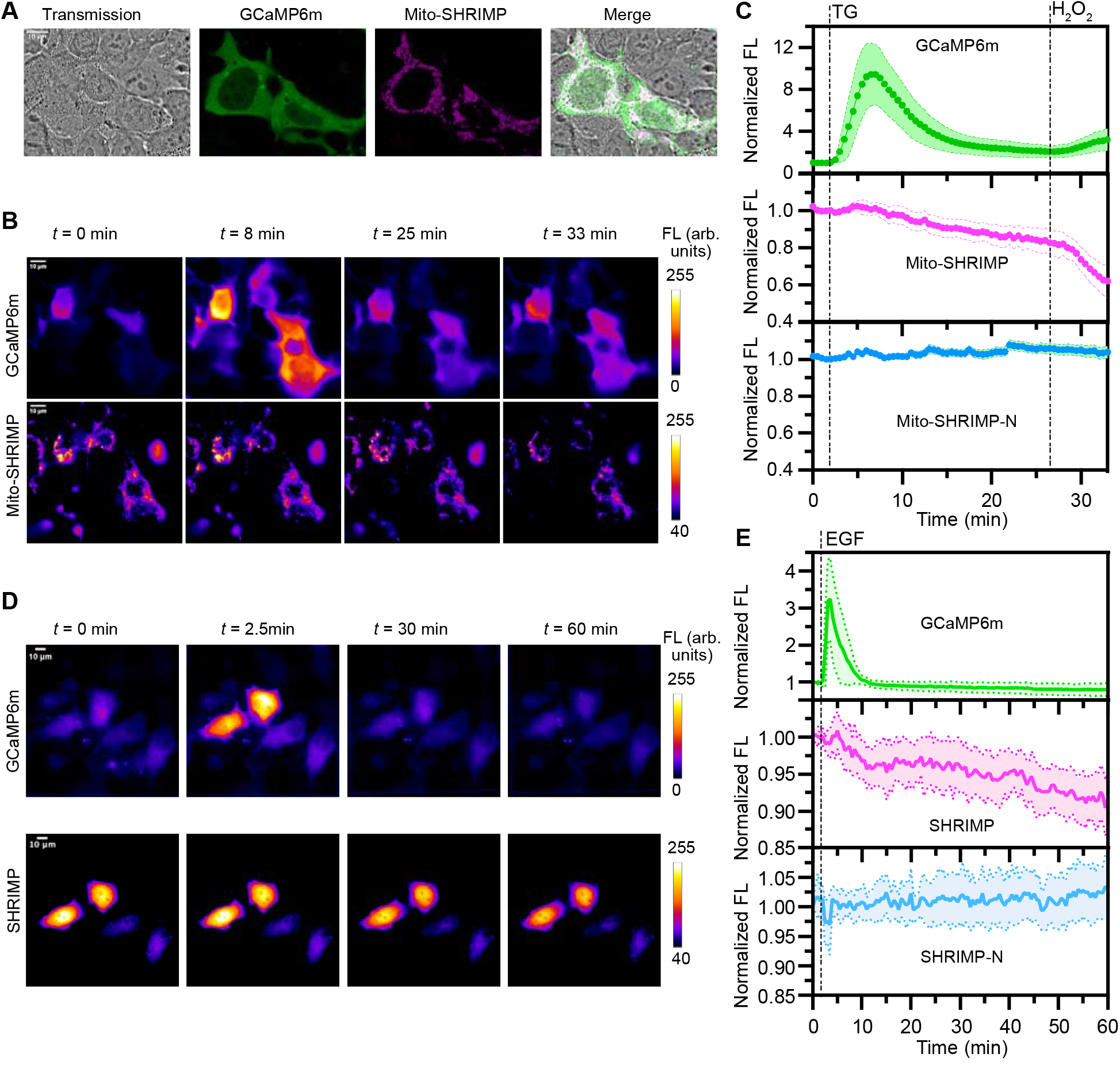
Combining SHRIMP with GCaMP6m for dual-color imaging of H_2_O_2_ and Ca^2+^ in mammalian cells. (A) Representative confocal fluorescence images of HEK 293T cells co-expressing GCaMP6m and mitochondrial (Mito-) SHRIMP. Scale bar: 10 μm. (B) Representative pseudocolored timelapse wide-field fluorescence images of HEK 293T cells co-expressing GCaMP6m and Mito-SHRIMP in response to sequential thapsigargin (TG, 10 μM) and H_2_O_2_ (150 μM) treatments. Scale bar: 10 μm. (C) Quantitation of timelapse responses of GCaMP6m (top), Mito-SHRIMP (middle), and Mito-SHRIMP-N (bottom) in HEK 293T cells. Data represent mean ± s.d. of 20 cells in each group from three technical repeats. (D) Representative pseudocolored timelapse wide-field fluorescence images of HeLa cells co-expressing GCaMP6m and SHRIMP in response to EGF treatment. Scale bar: 10 μm. (E) Quantitation of timelapse responses of GCaMP6m (top), SHRIMP (middle), and SHRIMP-N (bottom) in HeLa cells. Data represent mean ± s.d. of 27 cells in each group from three technical repeats. FL, fluorescence. Arb. units, arbitrary units.

In addition to HEK 293T cells, we also co-expressed SHRIMP and GCaMP6m in human cervical cancer HeLa cells. These cells were subjected to serum starvation followed by epidermal growth factor (EGF) stimulation. The treatment resulted in a robust increase in intracellular Ca^2+^, as evidenced by the drastic increase in GCaMP6m fluorescence (**Fig. 4DE**). Similar to the TG treatment, the rise and subsequent decay of the Ca^2+^ concentration occurred throughout a period of approximately 10 min, while the red fluorescence of SHRIMP gradually decreased and persisted throughout the nearly 1-hour monitoring period, indicating sustained H_2_O_2_ production. To further validate our observations, we expressed the negative control SHRIMP-N in both HEK 293T and HeLa cells. In these cells, we observed no response to TG, H_2_O_2_, or EGF treatments (**Fig. 4CE**). Collectively, our results confirm the specificity and reliability of SHRIMP for multicolor/multiparameter imaging when used in conjunction with other color-compatible GEFIs.

### Imaging of H_2_O_2_ in Primary Mouse Pancreatic Islets

Pancreatic islets are small clusters of cells found in the pancreas, with β-cells being responsible for insulin production and blood sugar regulation (56). However, oxidative stress can disrupt the function and integrity of these β-cells, contributing to the development of diabetes (57). Islet transplantation, a novel treatment for type 1 diabetes, involves the transplantation of functional islets from deceased donors into recipients (58,59). This procedure restores insulin production and improves glucose control. However, transplanted islets also face the challenge of oxidative stress, which can lead to islet death and reduced graft survival (60). To address this issue, ongoing research focuses on optimizing islet isolation techniques, improving transplant outcomes, and developing strategies to mitigate oxidative stress and protect transplanted islets from oxidative damage. In this context, there is a pressing need for innovative methods that can effectively monitor and assess oxidative stress levels in pancreatic islets.

To assess the reliability of SHRIMP in detecting H_2_O_2_ in pancreatic islets, we created adeno-associated viruses (AAV) containing the SHRIMP gene driven by the insulin promoter (pINS) to enable expression in β-cells. The AAV8 capsid was employed to optimize the virus for efficient transduction of islets (61). We effectively transduced primary mouse islets using this virus and achieved the expression of SHRIMP, as evidenced by the presence of bright red fluorescence. We next validated the responsiveness of SHRIMP in islets by directly adding H_2_O_2_. 5 μM of exogenous H_2_O_2_ did not trigger a notable response, while 25 and 100 μM H_2_O_2_ caused obvious fluorescence changes of SHRIMP. In parallel, 100 μM H_2_O_2_ did not affect the fluorescence of SHRIMP-N expressed in islets.

Streptozotocin (STZ) is a glucose analog that selectively accumulates in β-cells through glucose transporter 2 (GluT2) (62). It is known for its specific toxicity to pancreatic β-cells and has been utilized as a pancreatic cancer drug as well as a means to induce type 1 diabetes in animal models for research purposes. Although the precise molecular mechanism underlying STZ-induced β-cell toxicity remains incompletely understood, previous findings suggest that DNA alkylation and oxidative stress play key roles in this process (63,64). Enzymatic assays conducted in the past have demonstrated that millimolar concentrations of STZ can induce the production of H_2_O_2_ in isolated islets (64). Hence, we investigated the response of SHRIMP-expressing islets to STZ. Upon exposure to STZ at a concentration of 6 mM, there was a reproducible ∼30% reduction in SHRIMP fluorescence within 1 hour (**Fig. 5**). In parallel experiments, pancreatic islets expressing the negative control, SHRIMP-N, exhibited only a minor decrease in fluorescence intensity, which was likely attributed to photobleaching and pH alterations during β-cell damage. Nevertheless, these results provide support for the capability of SHRIMP to effectively monitor H_2_O_2_ production in primary islet tissue.

**Figure 5.**
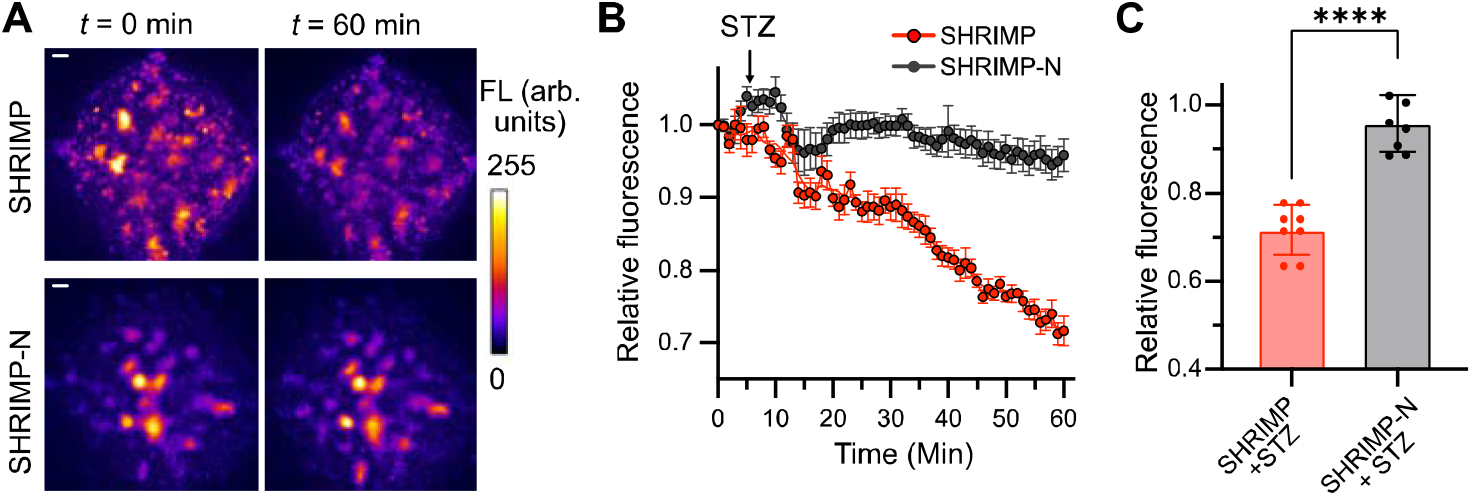
Detection of streptozotocin (STZ) mediated H_2_O_2_ in primary mouse pancreatic islet tissue. (A) Representative pseudocolored fluorescence images of islets expressing SHRIMP or SHRIMP-N. Scale bar: 10 μm. (B) Quantitation of timelapse responses. Arrow indicates the time point for STZ (6 mM) addition. (C) Comparison of fluorescence intensity changes at *t* = 60 min (\*\*\*\**P*<0.0001, unpaired two-tailed t-test). Data represent the mean and s.e.m. of eight randomly selected regions from three individual islets in three repeats. FL, fluorescence. Arb. units, arbitrary units.

## Discussion

In this study, we developed a circularly permutated red fluorescent protein, cpmScarlet, which exhibited significantly brighter red fluorescence compared to the cpmApple variant (**Table 1**). We fused cpmScarlet with the H_2_O_2_ sensory OxyR fragments, similar to the configurations of HyPerRed and other green fluorescent HyPer sensors (25,39). Through further optimization and mutagenesis, we identified SHRIMP, a genetically encoded red fluorescent H_2_O_2_ indicator with high brightness, specificity, resistance to blue-light-induced photoactivation, and a five-fold turn-off response to nanomolar H_2_O_2_.

A side-by-side comparison with HyPerRed demonstrated that SHRIMP outperformed it in terms of brightness and resistance to photoactivation (**Fig. 2**). While turn-on responses are generally preferred in redox biosensors, SHRIMP’s large turn-off response still provides a good dynamic range for monitoring H_2_O_2_ levels. Additionally, SHRIMP did not display detectable photoactivation, whereas HyPerRed showed notable photoactivation behavior. These findings highlight the superior characteristics of SHRIMP as a robust and reliable tool for studying H_2_O_2_ dynamics in living systems, making it a preferable choice over HyPerRed for redox imaging applications.

GEFIs based on cpmApple commonly suffer from low brightness and are susceptible to noticeable photoactivation when exposed to violet and blue light, which can introduce experimental artifacts (8,10,17). Although reducing blue light intensity may minimize these effects, our study offers an alternative that overcomes this limitation and can lead to more robust imaging tools. The development of cpmScarlet and SHRIMP, along with the fact that SHRIMP inherited excellent photophysical properties of mScarlet, strongly suggests that cpmScarlet is a preferable scaffold for developing future red GEFIs.

In addition to single FP-based indicators, cpmScarlet may be incorporated into fluorescent indicators based on Föster resonance energy transfer (FRET), serving as either a donor or acceptor. It should allow the creation of fusions in a different topology, compared to the original mScarlet, while the relative dipole orientation of the donor and acceptor is an important factor to consider when building FRET-based sensors (65).

We demonstrated the utility of SHRIMP to image H_2_O_2_ in live mammalian cells. SHRIMP robustly responded to exogenous H_2_O_2_ addition to HEK 293T cells and endogenously generated H_2_O_2_ in activated macrophages (**Fig. 3**). Furthermore, we combined SHRIMP with the calcium sensor GCaMP6m for dual-color dual-parameter imaging (**Fig. 4**). Our findings shed light on the intricate relationship between Ca^2+^ and H_2_O_2_ signals. The simultaneous monitoring of H_2_O_2_ levels and Ca^2+^ dynamics using SHRIMP and GCaMP6m not only demonstrates the versatility of SHRIMP in multicolor imaging but also opens up new avenues for investigating the interplay between ROS and Ca^2+^ signaling in various biological processes.

Moreover, we successfully employed SHRIMP to visualize H_2_O_2_ production in damaged pancreatic islet tissue. Our findings demonstrate the potential of SHRIMP for monitoring the redox dynamic in islets, which may advance our understanding of β-cell death and further improve cell replacement therapies for type 1 diabetes.

Recently, a more enhanced mScarlet mutant, mScarlet3, has been reported (17). We anticipate that these new mutations in mScarlet3 can be incorporated into cpmScarlet to further improve its robustness. Additionally, an OxyR domain from *Neisseria meningitidis*, which exhibits increased sensitivity to H_2_O_2_, has been utilized to develop a green GEFI called HyPer7 (41). In future work, it is possible to combine the new cpmScarlet mutant with the more sensitive OxyR domain to create even more advanced red fluorescent H_2_O_2_ indicators. This approach holds promise for further enhancing the sensitivity and performance of these biosensors in detecting and monitoring H_2_O_2_ dynamics.

In summary, our study showcases the feasibility of developing red GEFIs based on cpmScarlet and highlights the effectiveness of SHRIMP as a red GEFI for H_2_O_2_. SHRIMP offers several advantages, including enhanced brightness, resistance to photoactivation, and reliable responses both in purified protein form and within cells and tissues. This research sets the foundation for further advancements in red GEFIs and establishes SHRIMP as a valuable tool for investigating the dynamics of H_2_O_2_ in living systems.

## Materials and Methods

### Key Materials and General Methods

Chemicals were procured from Thermo Fisher Scientific or Sigma-Aldrich unless otherwise specified. DNA oligos were obtained from Eurofins Genomics or Integrated DNA Technologies. DNA sequencing services were provided by Eurofins Genomics. Phusion High-Fidelity DNA polymerase and restriction enzymes were purchased from Thermo Fisher Scientific. Taq DNA polymerase was acquired from New England Biolabs. Additional specific reagents used in this study include TG (Cat. # 10522) from Cayman Chemical, STZ (Cat. # ALX-380-010) from Enzo Life Sciences, and PMA (Cat. # P1585) and EGF (Cat. # GF316) from MilliporeSigma. The plasmids pCytERM_mScarlet_N1 (Addgene Plasmid # 85066), pC1-HyPer-Red (Addgene Plasmid # 48249), and pGP-CMV-GCaMP6m were generous gifts from Dorus Gadella, Vsevolod Belousov, and Douglas Kim (GENIE Project), respectively (25,42,66). C57BL/6J (The Jackson Laboratory, Cat. # 000664) wild-type mice were utilized for pancreatic islet isolation. The mice were housed in a temperature-controlled vivarium at ∼ 23°C and ∼ 50% humidity, with a 12-hour light-dark cycle. All animal procedures were conducted following the approved protocols by the Animal Care and Use Committees at the University of Virginia. ChatGPT was used to reword sentences and polish the manuscript.

### Protein Engineering and Library Screening

To obtain cpmScarlet, gene fragments encoding residues 149-232 and residues 2-146 of mScarlet were amplified from the pCytERM_mScarlet_N1 plasmid using Taq DNA Polymerase. The amplification was carried out using the oligos cpmScarlet-2NNK-F1 and cpmScarlet-R1, or cpmScarlet-F2 and cpmScarlet-NNK-R2 for the respective fragments. Subsequently, the two fragments were assembled through an overlap polymerase chain reaction (PCR) using the oligos cpmScarlet-2NNK-G-F and cpmScarlet-NNK-G-R. The resulting overlap PCR product was digested with Xho I and Hind III restriction enzymes and ligated into a predigested pBAD/His B plasmid. *E. coli* DH10B cells were transformed with the ligation product and plated on Luria-Bertani (LB) agar plates supplemented with 100 μg/mL ampicillin and 0.02% (w/v) L-arabinose. After incubation at 37°C for 24 hours, colonies exhibiting red fluorescence were screened using a previously described bacterial colony imaging system (67). Colonies with high brightness were selected to inoculate 0.5 mL of 2× YT medium supplemented with 100 μg/mL ampicillin and 0.02% (w/v) L-arabinose in a 96-well deep-well culture plate. The cells were initially grown at 250 rpm and 37°C until the optical density at 600 nm (OD_600_) reached 0.8, followed by further incubation at 250 rpm and 16 °C for an additional 48 hours. The cells were then pelleted by centrifugation, and the pellets were lysed with 300 μL of Bacterial Protein Extraction Reagents (B-PER, Pierce) in each well. The fluorescent intensity of each cell lysate was measured using a BioTek Synergy Mx Microplate Reader. Mutants exhibiting high-intensity red emission were selected for protein purification and further confirmation, ultimately leading to the identification of the final cpmScarlet mutant.

To engineer SHRIMP from cpmScarlet, the cpmScarlet gene was amplified from the corresponding pBAD plasmid using the oligos OxyR-cpmScarlet-F1 and OxyR-cpmScarlet-R1. The two OxyR fragments were amplified from pC1-HyPer-Red using the oligo pairs OxyR-N-F and OxyR-N-R, and OxyR-C-F and OxyR-C-R, respectively. The resulting products were assembled and inserted into a compatible pBAD/His B vector using Gibson assembly (68). The subsequent steps for library screening were similar to those described above for SHRIMP, with the exception that protein expression was induced at 16 °C and H_2_O_2_ (500 μM) was used to treat the prepared cell lysates. The fluorescent intensity of each cell lysate was recorded in the presence and absence of H_2_O_2_. The variant showing the maximal response, designated as SHRIMP0.1, was selected as the template for further directed evolution. Two rounds of random mutagenesis were performed using Taq DNA polymerase with oligos pBAD-F and pBAD-R by following a previously described procedure (67). Protein expression for these two libraries was induced at room temperature and 30°C, respectively. SHRIMP, which showed high brightness and extreme responsiveness, was identified from library screening.

### Protein Purification and In Vitro Characterization

Oligos pET-F and pET-R were used to amplify the SHRIMP gene from the corresponding pBAD plasmid and inserted into a pET-28a vector via Gibson assembly (68), maintaining the same reading frame with an N-terminal His_6_ tag. Protein expression was carried out using BL21(DE) cells at a temperature of 16 °C. The purification procedure, which involved sequential use of Ni-NTA agarose beads (Pierce) and a Cytiva HiLoad 16/60 Superdex 200 pg size-exclusion column, was conducted as previously described (67). Subsequently, the protein in phosphate-buffer saline (PBS, pH 7.4) was concentrated using an Amicon Ultra Centrifugal Filter Unit with a molecular weight cutoff of 10,000 Da. The concentrated protein was then divided into small aliquots and stored at −80 °C for further use. cpmScarlet, ecpApple, and HyPerRed proteins were prepared using similar procedures. Protein concentrations were determined using the alkaline denaturation method (67). Proteins were further diluted with PBS to a final concentration of 500 nM for recording excitation and emission spectra. Extinction coefficients and quantum yields were determined following a previous procedure (1), and mScarlet-I (Φ = 0.54) was used as the quantum yield reference (42). To examine the response of SHRIMP, the fully reduced protein was first prepared by mixing the protein (10 μM) with dithiothreitol (DTT, 1 mM). After 20 min incubation at room temperature, the mixture was diluted 1000-fold using PBS and then treated with oxidants. Fluorescence measurement was performed after 15 min of incubation on a BioTek Synergy Mx Microplate Reader with the excitation wavelength at 570 nm and the emission wavelength at 600 nm. pH titration was also determined according to a previously described procedure (67).

### Mammalian Cell Culture, Live Cell Imaging, and Flow Cytometry

To construct SHRIMP into mammalian cell expression vector pcDNA3, oligos pcDNA3-SHRIMP-F and pcDNA3-SHRIMP-R were used to amplify the SHRIMP insert from the pET-SHRIMP plasmid. The amplified fragment was inserted into a predigested pcDNA3 vector via Gibson assembly (68). Similar methods were used to generate the pcDNA3-SHRIMP-N and pcDNA3-HyPerRed plasmids. To construct a plasmid (pCS-MLS-SHRIMP) for mitochondrial localization of SHRIMP, oligos pMLS-F and pMLS-R were employed to amplify SHRIMP from the pcDNA3-SHRIMP plasmid. The amplified fragment was then cloned into a compatible pCS2+ vector (69) predigested with Nhe I and Xho I.

HEK 293T cells were cultured and transfected as previously described (67). Imaging was performed 24 hours post-transfection in Dulbecco’s PBS supplemented with 1 mM Ca^2+^ and 1 mM Mg^2+^. RAW 264.7 cells were cultured in Dulbecco’s Modified Eagle Medium (DMEM) supplemented with 10% FBS at 37 °C with 5% CO_2_. Transfection was performed with the X-tremeGENE HP DNA Transfection Reagent (Roche) according to the manufacturer’s instructions. RAW 264.7 cells were imaged in Hank’s Balanced Salt Solution (HBSS) supplemented with 1 mM Ca^2+^ and 1 mM Mg^2+^ at 24 hours post-transfection. A Leica DMi8 inverted microscope equipped with a Leica EL6000 light source, a Photometrics Prime 95B sCMOS camera, and a TRITC filter cube with 545/25-nm bandpass excitation and 605/70-nm bandpass emission was used for wide-field imaging. During time-lapse imaging, H_2_O_2_ (150 μM) and PMA (2 μM) were added to imaging buffers to stimulate HEK 293T and RAW 264.7 cells, respectively. Time-lapse images were analyzed as previously described (29).

To achieve dual-color imaging, the plasmids pGP-CMV-GCaMP6m and pCS-MLS-SHRIMP were used to co-transfect HEK 293T cells. Imaging was performed in the HBSS buffer supplemented with 1 mM Ca^2+^ and 1 mM Mg^2+^ at 48 hours post-transfection. To examine the subcellular localization, a Leica SPE-II spectral confocal module was used for confocal imaging. Green fluorescence was obtained using a 488 nm laser, and the emission was collected within the range of 510 to 565 nm. Red fluorescence, on the other hand, was acquired using a 532 nm laser, and the emission was collected within the range of 600 to 665 nm. For time-lapse imaging, the wide-field microscope system mentioned earlier was utilized. To collect the emission of GCaMP6m, an additional FITC filter cube with 470/40-nm bandpass excitation and 525/50-nm bandpass emission was employed. 10 μM TG was used to stimulate cells. 150 μM H_2_O_2_ was added near the end of the imaging session to further confirm the responsiveness of the sensors.

HeLa cells were cultured in DMEM supplemented with 15% FBS at 37 °C with 5% CO_2_. pGP-CMV-GCaMP6m and pcDNA-SHRIMP were used to transfect the cells as described (12). 48 hours after transfection, the cells were subjected to overnight serum starvation in DMEM containing only 0.1% FBS. Subsequently, the cells were rinsed three times with a mammalian cell imaging buffer (137 mM NaCl, 2.2 mM KCl, 2 mM CaCl_2_, 2 mM MgCl_2_, 2 mM glucose, 25 mM HEPES, pH 7.4) before being equilibrated in the imaging buffer for 30 min. The cells were then imaged using the aforementioned wide-field microscope setup. To stimulate the cells, EGF at a concentration of 200 ng/mL was used.

To compare the brightness of SHRIMP and HyPerRed in HEK 293T cells, flow cytometry analysis was conducted 24 hours after transfection. Cells were resuspended in a buffer (pH 7.4) containing 145 mM NaCl, 5 mM KCl,1.2 mM MgCl_2_, 2.6 mM CaCl_2_, 10 mM HEPES, and 1% FBS. An Attune NxT Flow Cytometer (Thermo Fisher) was utilized to analyze more than 20,000 cells for each group. The emission was set to 561 nm, and the red fluorescence signal was collected using the YL2 channel with a 620/15 bandpass filter.

### Characterization of Photostability and Photoactivation

For photostability and photoactivation characterization, purified SHRIMP or HyPerRed proteins were diluted to a concentration of 100 μM in PBS. To prepare the experimental setup, 500 μL of Remel mineral oil (Thermo Scientific, Cat. # R21237) was washed twice with an equal volume of ddH_2_O, followed by another wash with an equal volume of PBS. The mineral oil was then equilibrated with an equal volume of PBS. Next, 1 μL of the diluted protein was mixed with 50 μL of the pre-treated mineral oil by vortexing, resulting in the formation of small protein-oil droplets. These droplets were loaded onto the bottom of a 35-mm diameter glass bottom dish. For photobleaching comparison, the protein droplets were subjected to continuous wide-field illumination using the aforementioned DMi8 microscope equipped with the TRITC filter set and a 40× oil objective (NA 1.15). Two different excitation light intensities were used: 10% and 100%. The optical power was measured using a digital optical power meter (Thorlabs, #PM100D) with a microscope slide power meter sensor (Thorlabs, #S170C). The light intensity at the focal plane was approximately 0.09 and 1 W/cm^2^, respectively. To examine the photoactivation behavior, the Leica SPE-II spectral confocal module was employed. A 275 μm × 275 μm area was scanned five times using a 532 nm laser set at 50% power (∼366.8 μW at the focal plane, as determined by the power meter), followed by one scan using a 405 nm laser at 80% power intensity (∼558.7 μW at the focal plane, as determined by the power meter). The scanning was performed at 400 Hz with 512×512 pixels, and the pixel dwell time was set to 1.44 μs. To analyze imaging data, random regions of interest (ROIs) were selected, and the mean fluorescence intensity at specific times (*F*_t_) was normalized to the mean fluorescence intensity of the first frame (*F*_0_), and the ratio (*F*_t_/*F*_0_) was plotted against time.

### Viral Preparation, and Transduction and Imaging of Mouse Pancreatic Islets

The SHRIMP or SHRIMP-N gene was amplified with oligos pAAV-insulin-F and pAAV-insulin-R from the corresponding pcDNA3 mammalian cell expression plasmids and inserted into a modified pAAV vector harboring a human insulin promoter (pINS) via Gibson Assembly. AAV packing, purification, and quantitative PCR (qPCR) titer quantification were conducted following the previously described protocol (70), with the only difference being the use of the pAAV2/8 packing plasmid (Addgene Plasmid #112864) for serotyping. The resulting titers for SHRIMP and SHRIMP-N AAVs were determined to be 2×10^12^ and 1×10^13^ genome copies per mL, respectively. The AAVs were aliquoted and stored at −80 °C. Mouse pancreatic islets were isolated and cultured as previously described (71). For AAV transduction, islets were pre-treated with 0.05% trypsin at 37 °C for 2 min and immediately transferred into Roswell Park Memorial Institute (RPMI) 1640 Medium supplemented with 10% fetal bovine serum (FBS) to stop trypsinization. About 50 islets were selected and placed into each well of a 96-well plate containing 100 μL of RPMI 1640 Medium supplemented with 10% FBS. Next, 5 μL of the stock AAV was added to each well. The plate was then incubated at 37 °C with 5% CO_2_ for 20 hours. Following the incubation period, both the cells and the medium were transferred from the 96-well plates to 24-well plates, with each well containing 1 mL of RPMI 1640 Medium supplemented with 10% FBS. The medium was replaced with fresh RPMI 1640 Medium supplemented with 10% FBS after 2 days, and the islets were imaged one day following the medium change. Before imaging, islets were transferred into a Krebs-Ringer Modified Buffer (135 mM NaCl, 5 mM KCl, 1 mM MgSO_4_, 0.4 mM K_2_HPO_4_, 1 mM CaCl_2_, 5.5 mM Glucose, and 20 mM HEPES, pH 7.4). H_2_O_2_ (from 5 to 100 μM) or STZ (6 mM) was added to the imaging buffer. Images were acquired using the abovementioned wide-field condition.

## Acknowledgments

Research reported in this publication was supported by the University of Virginia, the National Institutes of Health under grants R01GM129291, R01DK122253, and RF1AG077773, and the United States-Israel Binational Science Foundation (BSF) under grant 2021256.

## Author Contributions

H.-w.A. conceived and supervised the whole project. Y.P. and Y.Z. developed and characterized the SHRIMP indicator with contributions from J.Z. and Z.L. Y.H. prepared primary mouse islets and Y.W. and J.O. supervised Y.H.’s experiments. H.-w.A., Y.P., and Y.Z. wrote the manuscript with inputs from other authors.

## References

1. E. C. Greenwald, S. Mehta, J. Zhang, Genetically Encoded Fluorescent Biosensors Illuminate the Spatiotemporal Regulation of Signaling Networks. Chem. Rev. 118, 11707–11794 (2018).

2. Z. Chen, T. M. Truong, H. Ai, Illuminating Brain Activities with Fluorescent Protein-Based Biosensors. Chemosensors 5, 32 (2017).

3. Y. Shen, T. Lai, R. E. Campbell, Red fluorescent proteins (RFPs) and RFP-based biosensors for neuronal imaging applications. Neurophotonics 2, 031203 (2015).

4. T. Wu et al., A Genetically Encoded Far-Red Fluorescent Indicator for Imaging Synaptically-Released Zn^2+^. Sci. Adv. 9, eadd2058 (2023).

5. G. S. Baird, D. A. Zacharias, R. Y. Tsien, Circular permutation and receptor insertion within green fluorescent proteins. Proc. Natl. Acad. Sci. U.S.A. 96, 11241–11246 (1999).

6. Y. Nasu, Y. Shen, L. Kramer, R. E. Campbell, Structure- and mechanism-guided design of single fluorescent protein-based biosensors. Nat. Chem. Biol. 17, 509–518 (2021).

7. Y. Zhao et al., An expanded palette of genetically encoded Ca^2+^ indicators. Science 333, 1888–1891 (2011).

8. J. Akerboom et al., Genetically encoded calcium indicators for multi-color neural activity imaging and combination with optogenetics. Front. Mol. Neurosci. 6, 2 (2013).

9. H. J. Carlson, R. E. Campbell, Circular permutated red fluorescent proteins and calcium ion indicators based on mCherry. Protein Eng. Des. Sel. 26, 763–772 (2013).

10. Y. Shen et al., A genetically encoded Ca^2+^ indicator based on circularly permutated sea anemone red fluorescent protein eqFP578. BMC Biol. 16, 9 (2018).

11. T. Patriarchi et al., An expanded palette of dopamine sensors for multiplex imaging in vivo. Nat. Methods 17, 1147–1155 (2020).

12. Y. Fan, M. Makar, M. X. Wang, H. W. Ai, Monitoring thioredoxin redox with a genetically encoded red fluorescent biosensor. Nat. Chem. Biol. 13, 1045–1052 (2017).

13. Y. Ohta, T. Furuta, T. Nagai, K. Horikawa, Red fluorescent cAMP indicator with increased affinity and expanded dynamic range. Sci. Rep. 8, 1866 (2018).

14. Y. Pang, H. Zhang, H. W. Ai, Genetically Encoded Fluorescent Redox Indicators for Unveiling Redox Signaling and Oxidative Toxicity. Chem. Res. Toxicol. 34, 1826–1845 (2021).

15. Z. Chen, H. W. Ai, Single Fluorescent Protein-Based Indicators for Zinc Ion (Zn^2+^). Anal. Chem. 88, 9029–9036 (2016).

16. Q. Ni, S. Mehta, J. Zhang, Live-cell imaging of cell signaling using genetically encoded fluorescent reporters. FEBS J. 285, 203–219 (2018).

17. T. W. J. Gadella, Jr. et al., mScarlet3: a brilliant and fast-maturing red fluorescent protein. Nat. Methods 20, 541–545 (2023).

18. J. Zuo et al., Redox signaling at the crossroads of human health and disease. MedComm 3, e127 (2022).

19. B. C. Dickinson, C. J. Chang, Chemistry and biology of reactive oxygen species in signaling or stress responses. Nat. Chem. Biol. 7, 504–511 (2011).

20. M. Valko, C. Rhodes, J. Moncol, M. Izakovic, M. Mazur, Free radicals, metals and antioxidants in oxidative stress-induced cancer. Chem. Biol. Interact. 160, 1–40 (2006).

21. K. J. Barnham, C. L. Masters, A. I. Bush, Neurodegenerative diseases and oxidative stress. Nat. Rev. Drug Discov. 3, 205–214 (2004).

22. I. Liguori et al., Oxidative stress, aging, and diseases. Clin. Interv. Aging 13, 757 (2018).

23. A. I. Kostyuk et al., In Vivo Imaging with Genetically Encoded Redox Biosensors. Int. J. Mol. Sci. 21 (2020).

24. M. Schwarzlander, T. P. Dick, A. J. Meyer, B. Morgan, Dissecting Redox Biology Using Fluorescent Protein Sensors. Antioxid. Redox Signal. 24, 680–712 (2016).

25. Y. G. Ermakova et al., Red fluorescent genetically encoded indicator for intracellular hydrogen peroxide. Nat. Commun. 5, 5222 (2014).

26. A. G. Shokhina et al., Red fluorescent redox-sensitive biosensor Grx1-roCherry. Redox Biol. 21, 101071 (2018).

27. Y. Fan, Z. Chen, H. W. Ai, Monitoring redox dynamics in living cells with a redox-sensitive red fluorescent protein. Anal. Chem. 87, 2802–2810 (2015).

28. Y. Pang, H. Zhang, H. W. Ai, Improved Red Fluorescent Redox Indicators for Monitoring Cytosolic and Mitochondrial Thioredoxin Redox Dynamics. Biochemistry 61, 377–384 (2022).

29. Y. Pang et al., Development, Characterization, and Structural Analysis of a Genetically Encoded Red Fluorescent Peroxynitrite Biosensor. ACS Chem. Biol. 18, 1388–1397 (2023).

30. M. P. Murphy, How mitochondria produce reactive oxygen species. Biochem. J. 417, 1–13 (2009).

31. J. Zielonka et al., Recent Developments in the Probes and Assays for Measurement of the Activity of NADPH Oxidases. Cell Biochem. Biophys. 75, 335–349 (2017).

32. D. R. Gough, T. G. Cotter, Hydrogen peroxide: a Jekyll and Hyde signalling molecule. Cell Death Dis. 2, e213 (2011).

33. H. Antelmann, J. D. Helmann, Thiol-based redox switches and gene regulation. Antioxid. Redox Signal. 14, 1049–1063 (2011).

34. T. Tiganis, Reactive oxygen species and insulin resistance: the good, the bad and the ugly. Trends Pharmacol. Sci. 32, 82–89 (2011).

35. Y. S. Bae et al., Epidermal growth factor (EGF)-induced generation of hydrogen peroxide: role in EGF receptor-mediated tyrosine phosphorylation. J. Biol. Chem. 272, 217–221 (1997).

36. A. Corcoran, T. G. Cotter, Redox regulation of protein kinases. FEBS J. 280, 1944–1965 (2013).

37. M. Schieber, N. S. Chandel, ROS function in redox signaling and oxidative stress. Curr. Biol. 24, R453–R462 (2014).

38. K. M. Holmström, T. Finkel, Cellular mechanisms and physiological consequences of redox-dependent signalling. Nat. Rev. Mol. Cell Biol. 15, 411–421 (2014).

39. V. V. Belousov et al., Genetically encoded fluorescent indicator for intracellular hydrogen peroxide. Nat. Methods 3, 281–286 (2006).

40. D. S. Bilan et al., HyPer-3: a genetically encoded H_2_O_2_ probe with improved performance for ratiometric and fluorescence lifetime imaging. ACS Chem. Biol. 8, 535–542 (2013).

41. V. V. Pak et al., Ultrasensitive Genetically Encoded Indicator for Hydrogen Peroxide Identifies Roles for the Oxidant in Cell Migration and Mitochondrial Function. Cell Metab. 31, 642–653.e646 (2020).

42. D. S. Bindels et al., mScarlet: a bright monomeric red fluorescent protein for cellular imaging. Nat. Methods 14, 53–56 (2017).

43. J. Zhang, Z. Li, Y. Pang, Y. Fan, H. W. Ai, Genetically Encoded Boronolectin as a Specific Red Fluorescent UDP-GlcNAc Biosensor. ACS Sens, DOI: 10.1021/acssensors.1023c00409 (2023).

44. B. K. Huang, H. D. Sikes, Quantifying intracellular hydrogen peroxide perturbations in terms of concentration. Redox Biol. 2, 955–962 (2014).

45. M. Canton et al., Reactive Oxygen Species in Macrophages: Sources and Targets. Front. Immunol. 12, 734229 (2021).

46. O. Inanami et al., Activation of the leukocyte NADPH oxidase by phorbol ester requires the phosphorylation of p47PHOX on serine 303 or 304. J. Biol. Chem. 273, 9539–9543 (1998).

47. E. C. Larsen et al., Differential requirement for classic and novel PKC isoforms in respiratory burst and phagocytosis in RAW 264.7 cells. J. Immunol. 165, 2809–2817 (2000).

48. M. S. Giambelluca, O. A. Gende, Hydrogen peroxide activates calcium influx in human neutrophils. Mol. Cell Biochem. 309, 151–156 (2008).

49. Q. Hu, S. Corda, J. L. Zweier, M. C. Capogrossi, R. C. Ziegelstein, Hydrogen peroxide induces intracellular calcium oscillations in human aortic endothelial cells. Circulation 97, 268–275 (1998).

50. M. Trebak, R. Ginnan, H. A. Singer, D. Jourd’heuil, Interplay between calcium and reactive oxygen/nitrogen species: an essential paradigm for vascular smooth muscle signaling. Antioxid. Redox Signal. 12, 657–674 (2010).

51. R. Sakurada et al., Calcium Release from Endoplasmic Reticulum Involves Calmodulin-Mediated NADPH Oxidase-Derived Reactive Oxygen Species Production in Endothelial Cells. Int. J. Mol. Sci. 20 (2019).

52. K. T. Jones, G. R. Sharpe, Thapsigargin raises intracellular free calcium levels in human keratinocytes and inhibits the coordinated expression of differentiation markers. Exp. Cell Res. 210, 71–76 (1994).

53. O. Thastrup, P. J. Cullen, B. K. Drøbak, M. R. Hanley, A. P. Dawson, Thapsigargin, a tumor promoter, discharges intracellular Ca^2+^ stores by specific inhibition of the endoplasmic reticulum Ca^2+^-ATPase. Proc. Natl. Acad. Sci. U. S. A. 87, 2466–2470 (1990).

54. P. S. Herson, K. Lee, R. D. Pinnock, J. Hughes, M. L. Ashford, Hydrogen peroxide induces intracellular calcium overload by activation of a non-selective cation channel in an insulin-secreting cell line. J. Biol. Chem. 274, 833–841 (1999).

55. H. Sato et al., Hydrogen peroxide mobilizes Ca2+ through two distinct mechanisms in rat hepatocytes. Acta Pharmacol. Sin. 30, 78–89 (2009).

56. P. Rorsman, M. Braun, Regulation of Insulin Secretion in Human Pancreatic Islets. Annu. Rev. Physiol., Vol 75 75, 155–179 (2013).

57. N. Eguchi, N. D. Vaziri, D. C. Dafoe, H. Ichii, The Role of Oxidative Stress in Pancreatic β Cell Dysfunction in Diabetes. Int. J. Mol. Sci. 22 (2021).

58. A. M. Shapiro et al., Islet transplantation in seven patients with type 1 diabetes mellitus using a glucocorticoid-free immunosuppressive regimen. N. Engl. J. Med. 343, 230–238 (2000).

59. A. Gangemi et al., Islet transplantation for brittle type 1 diabetes: the UIC protocol. Am. J. Transplant. 8, 1250–1261 (2008).

60. J. M. Barra, H. M. Tse, Redox-Dependent Inflammation in Islet Transplantation Rejection. Front. Endocrinol. 9, 175 (2018).

61. A. Ramzy et al., AAV8 Ins1-Cre can produce efficient β-cell recombination but requires consideration of off-target effects. Sci. Rep. 10, 10518 (2020).

62. W. J. Schnedl, S. Ferber, J. H. Johnson, C. B. Newgard, STZ transport and cytotoxicity. Specific enhancement in GLUT2-expressing cells. Diabetes 43, 1326–1333 (1994).

63. S. P. LeDoux, S. E. Woodley, N. J. Patton, G. L. Wilson, Mechanisms of nitrosourea-induced beta-cell damage. Alterations in DNA. Diabetes 35, 866–872 (1986).

64. N. T. Friesen, A. S. Büchau, P. Schott-Ohly, A. Lgssiar, H. Gleichmann, Generation of hydrogen peroxide and failure of antioxidative responses in pancreatic islets of male C57BL/6 mice are associated with diabetes induced by multiple low doses of streptozotocin. Diabetologia 47, 676–685 (2004).

65. H. W. Ai, Fluorescent-protein-based probes: general principles and practices. Anal. Bioanal. Chem. 407, 9–15 (2015).

66. T. W. Chen et al., Ultrasensitive fluorescent proteins for imaging neuronal activity. Nature 499, 295–300 (2013).

67. T. Wu, Y. Pang, H. W. Ai, Circularly Permuted Far-Red Fluorescent Proteins. Biosensors 11, 438 (2021).

68. D. G. Gibson et al., Enzymatic assembly of DNA molecules up to several hundred kilobases. Nat. Methods 6, 343–345 (2009).

69. R. A. Rupp, L. Snider, H. Weintraub, Xenopus embryos regulate the nuclear localization of XMyoD. Genes Dev. 8, 1311–1323 (1994).

70. X. Tian et al., A luciferase prosubstrate and a red bioluminescent calcium indicator for imaging neuronal activity in mice. Nat. Commun. 13, 3967 (2022).

71. M. Chen et al., Genetically Encoded, Photostable Indicators to Image Dynamic Zn^2+^ Secretion of Pancreatic Islets. Anal. Chem. 91, 12212–12219 (2019).

